# *Ixodes ricinus* tick bacteriome alterations based on a climatically representative survey in Hungary

**DOI:** 10.1101/2022.10.15.512391

**Authors:** Adrienn Gréta Tóth, Róbert Farkas, Márton Papp, Oz Kilim, Haeun Yun, László Makrai, Gergely Maróti, Mónika Gyurkovszky, Eszter Krikó, Norbert Solymosi

## Abstract

**Background:** The microbial communities of disease vectors may represent a key feature in several biological functions and thus deserve special attention in light of climate change and the consequent need to develop novel control strategies. Nevertheless, vector-borne microbial networks are still poorly understood. Assessing vectors’ microbial interactions and climatic dependencies may contribute to better-estimating pathogen transmission characteristics, public health risks and the urgency for control steps to be taken.

**Methods:** In a climatically representative country-wide survey, *Ixodes ricinus* ticks were collected from 17 locations in Hungary. Using shotgun metagenomic sequencing, the bacteriome composition was analyzed by investigating the relationship between the abundances of nymphs and females in various climatic environments.

**Results:** Bacterial composition on the genus level revealed a significant difference between the samples from females and nymphs. Within the core bacteriome, females and nymphs showed significant variation in the following genera: *Arsenophonus, Bacillus, Candidatus Midichloria, Rhodococcus, Sphingomonas, Staphylococcus, Wolbachia*. Among females, according to temperature strata, the following were found differentiating: *Curtobacterium, Pseudomonas, Sphingomonas*. There was no genus with a significant difference in precipitation categories for females. In the nymphs, *Curtobacterium* showed significant variation between temperature and *Bacillus* and *Curtobacterium* for various precipitation levels. Based on the full sample set, *Arsenophonus* and *Wolbachia* correlated positively which we assumed to have occurred due to the presence of *I. hookeri*.

**Conclusions:** The composition of vector-borne bacteriome members showed significant alterations at sampling points with different climatic conditions and development stages of the tick hosts. Our findings not only pave the way towards understanding tick-borne bacterial networks and interdependencies but also shed light on the high potential for the presence of a possible biological tick control species, the tick parasitoid, *Ixodiphagus hookeri* based on related bacteriome patterns. The results of conscious tick microbiome assessment studies may contribute to precision tick control strategies of the future.

## Introduction

Almost all aspects of human life are directly or indirectly affected by the Earth’s climate system dynamics. Many organisms, such as arthropods being poikilotherm organisms, are particularly sensitive to climate factors.^1^ Recent estimates suggest that around 80% of the world’s human population is at risk to one or more vector-borne disease(s) (VBDs).^2^ The geographical distribution of arthropod vectors is changing due to climate change.^3,4^ The response of ticks and dipteran vectors to the changing climate appears to differ.^5^ While bivalves respond to short-term weather and climate changes with rapid responses, ticks are affected by spatio-temporal averages of climate variation rather than short term or localised climate variability. This suggests that changes in the risk of disease spread by dipterans can be expected in the short term, while the prevalence of tick-borne diseases (TBDs) may be expected over a more extended period of time.^5^ *Ixodes ricinus* is the primary vector of the most prevalent TBDs in Europe, tick-borne encephalitis (TBE) and Lyme borreliosis (LB). Although reports on the prevalence of these TBDs do not show consistent global trends,^6,7^ the European *I. ricinus* populations do. As a consequence of climate change, *I. ricinus* has been found at extreme altitudes and latitudes other than its former range and its population has shifted further north within the European continent.^4,8,9^ Similarly, the geographic range of LB has expanded, specifically towards higher altitudes and latitudes.^6,7^ In consideration of climatic factors, a shift towards a more thermophilic tick fauna has also been described in Hungary.^10^

The eukaryotic, and as such, tick-borne microbiota impacts numerous biological functions of the host causing a variety of detrimental, neutral, or beneficial effects. In addition, certain bacterial assemblages can affect tick-borne pathogens (TBPs) of public health importance. Thus, understanding the changes in tick microbiota associated with the climatic environment could aide in the deeper understanding of tick biology.^11,12^ Because the tick bacteriota contains a number of climate-dependent, environment-dependent components, bacterial interaction patterns may also be determined by climatic factors.^13^ In our work, we investigated the diversity of the bacteriome of *I. ricinus* samples from nymphs and females from wetter or drier and cooler or warmer environments based on a climatically representative survey in Hungary.

## Materials and Methods

### Sampling design and sample collection

As the study’s main goal was to understand the natural bacteriome differences in *I. ricinus*, we designed the sampling to be representative to Hungary. In order to achieve this, we identified sampling points representative of climatic conditions.^14^ In Hungary, there are 175 local administrative units (LAU 1), for each of these units (geographical areas), we calculated the 10-year average of the yearly growing degree days (GDD) with base 10°C and the total yearly precipitation for each of the years. Meteorology data for the period of 2008 to 2017 was gathered from the ERA-Interim reanalysis data repository^15^ with a spatial resolution of 0.125°. For each of the two environmental variables, binary categories were defined: GDD with classes “cooler” and “warmer” and precipitation with classes “less” and “more”. Regarding GDD, the lower two quartiles were classified as “cooler” and the upper two quartiles as “warmer”. For precipitation, the yearly means below the country-wide median was defined as “less” and scores above the median as “more”. Each LAU 1 was categorized with its own climatic variables. By stratified spatial random sampling,^16,17^ twenty local administrative units were chosen as sampling areas. The strata’s sample size was proportional to the stratifying GDD and precipitation categories’ country-wide frequency to be representative. All data management and analysis was performed in the R environment.^18^ A forest edge to sample from was identified within each of the 20 selected LAUs. Between 23/3/2019 and 20/5/2019, questing ticks were collected by flagging and dragging. In the laboratory, ticks from the frozen samples were classified taxonomically. Ten nymphs and ten females of *I. ricinus* were selected per sampling site. Since we could not collect the minimum of ten females and ten nymphs at three sampling sites, only the samples collected at the remaining 17 sites were included in the sequencing and downstream analyses (Fig 1). Before DNA extraction, the ticks were washed twice with 99.8% alcohol.

**Figure 1.**
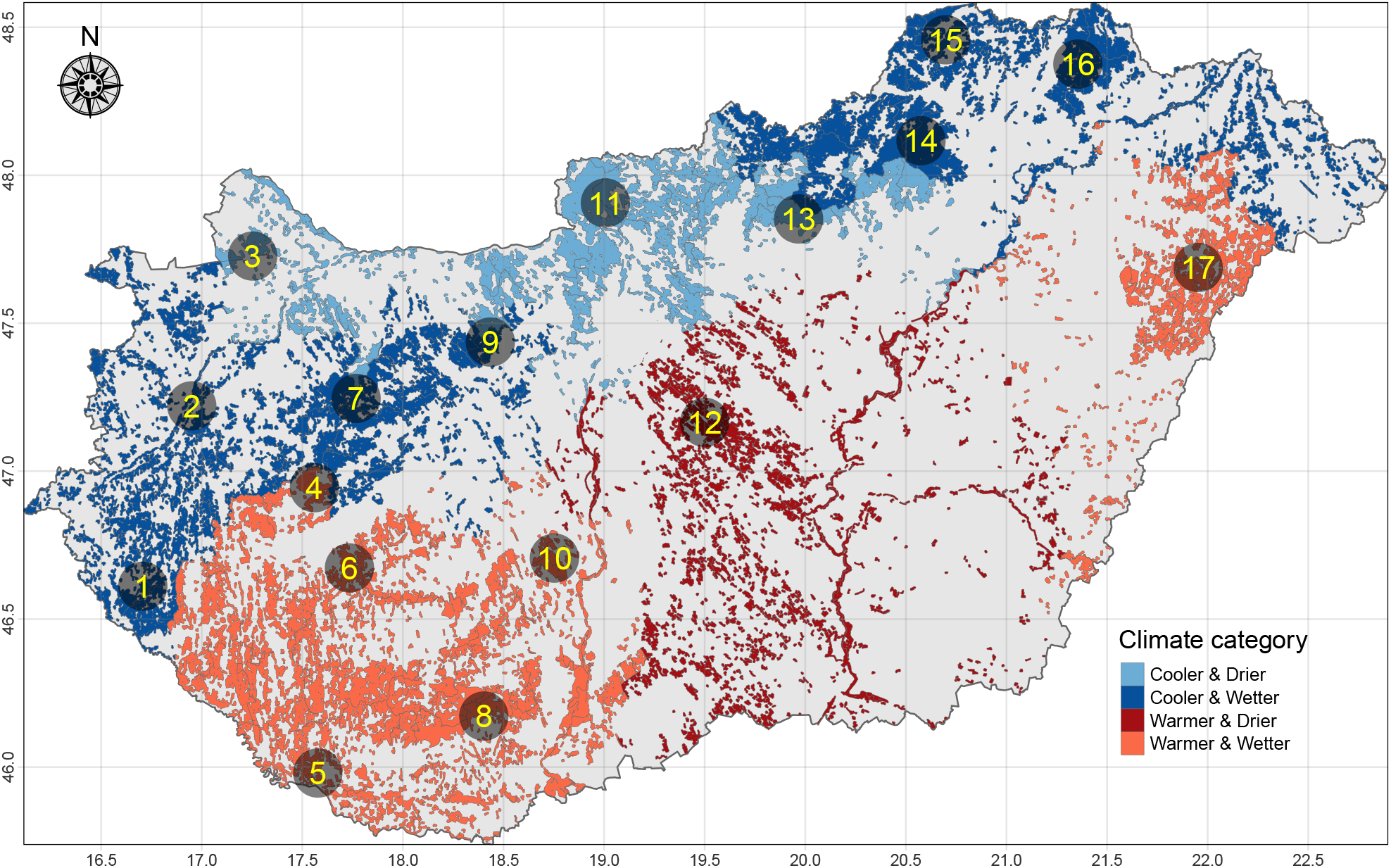
Climate category spatial pattern and sampling points. The color of the sylvan areas is defined by the climatic categories based on growing degree days (GDD) and precipitation over the period 2008-2017. The points are relatively homogeneous within Hungary geographically

### DNA extraction and metagenomics library preparation

The blackPREP Tick DNA/RNA Kit (Analytik Jena GmbH) was used for the DNA isolation. Isolated total metagenome DNA was used for library preparation. In vitro fragment libraries were prepared using the NEBNext Ultra II DNA Library Prep Kit for Illumina. Paired-end fragment reads were generated on an Illumina NextSeq sequencer using TG NextSeq 500/550 High Output Kit v2 (300 cycles). Primary data analysis (base-calling) was carried out with Bbcl2fastq software (v2.17.1.14, Illumina).

### Bioinformatic analysis

After merging the paired-end reads by PEAR^19^, quality-based filtering and trimming were performed by TrimGalore (v.0.6.6, https://github.com/FelixKrueger/TrimGalore), with a setting of 20 as a quality threshold, retaining reads longer than 50 bp only. The remaining reads after deduplication by VSEARCH^20^ were taxonomically classified using Kraken2 (k=35)^21^ with a database created (26/03/2022) from the NCBI RefSeq complete archaeal, bacterial, viral genomes. For taxon assignment the `-confidence 0.5` parameter was used to obtain more precise hits. Core bacteria was defined as the relative abundance of agglomerated counts on genus-level above 1% in at least one of the samples. The taxon classification data was managed in R^18^ using functions of package phyloseq,^22^ microbiome^23^ and metacoder.^24^ The pre-processed reads were assembled to contigs by MEGAHIT (v1.2.9)^25^ using default settings. The contigs were also classified taxonomically by Kraken2 with the same database as above. The assembly-generated contigs that were classified to a pathogen bacteria genus by Kraken2 we reclassified by BLAST^26^ on the representative prokaryotic genomes (downloaded on 16/6/2022). For each contig, the longest and the smallest e-value hit were kept and reported.

### Statistical analysis

The within-subject (*α*) diversity was assessed using the numbers of observed species (richness) and the Inverse Simpson’s Index (evenness). These indices were calculated in 1000 iterations of rarefied OTU tables with a sequencing depth of 158. The average over the iterations was taken for each sample. The *α*-diversity expressed by Inverse Simpson’s Index was compared between the conditions using linear models. Comparing the female and nymph samples collected, a mixed-effect model was applied to manage the repeated measurements by sampling site as a random factor.

The between-subject (*β*) diversity was assessed by UniFrac distance^27^ based on the relative abundances of bacteria species. Using this measure, principal coordinate analysis (PCoA) ordination was applied to visualise the samples’ variance. To examine statistically whether the bacterial species composition differed by strata PERMANOVA (Permutational Multivariate Analysis of Variance^28^) was performed using the package vegan^29^ in R.^18^

The abundance differences in core bacteriome between groups were analysed by a negative binomial generalised model of DESeq2 package^30^ in R.^18^ This approach was applied following the recommendation of Weiss et al.^31^ None of the compared groups had more than 20 samples, and their average library size ratio was less than 10. According to the multiple comparisons, the FDR-adjusted p-value less than 0.05 was considered significant. The SparCC correlation coefficient quantified the relationship among the relative abundances of bacterial species.^32,33^ The statistical tests were two-sided.

## Results

After the basic demography of the samples, we present the *α*-diversity of the full bacteriome. From the analysis of the core bacteriome, we report the species that are part of it, the *β* -diversity, and the differences of the genera level expressed relative abundances. The Supplementary Information section summarizes species of core bacteriome, the detected pathogen bacteria and the correlations of bacteria genera (Fig 6).

Among the ticks collected at the sampling sites, the median proportion of nymphs was 76.52% (IQR: 19.33), females 15.32% (IQR: 8.58) and males 7.91% (IQR: 11.47).

The numbers of observed species and the Inverse Simpson’s Index *α*-diversity metrics by strata are shown in Fig 2. Alpha diversity showed no significant difference between groups in either metric. For the comparison of females and nymphs, the p-value was 0.138. Within females, the p-values obtained by comparing groups from colder and warmer environments and from drier and wetter environments were p=0.562 and p=0.577, respectively. Within nymphs, the comparative metrics were p=0.174 and p=0.309, respectively.

**Figure 2.**
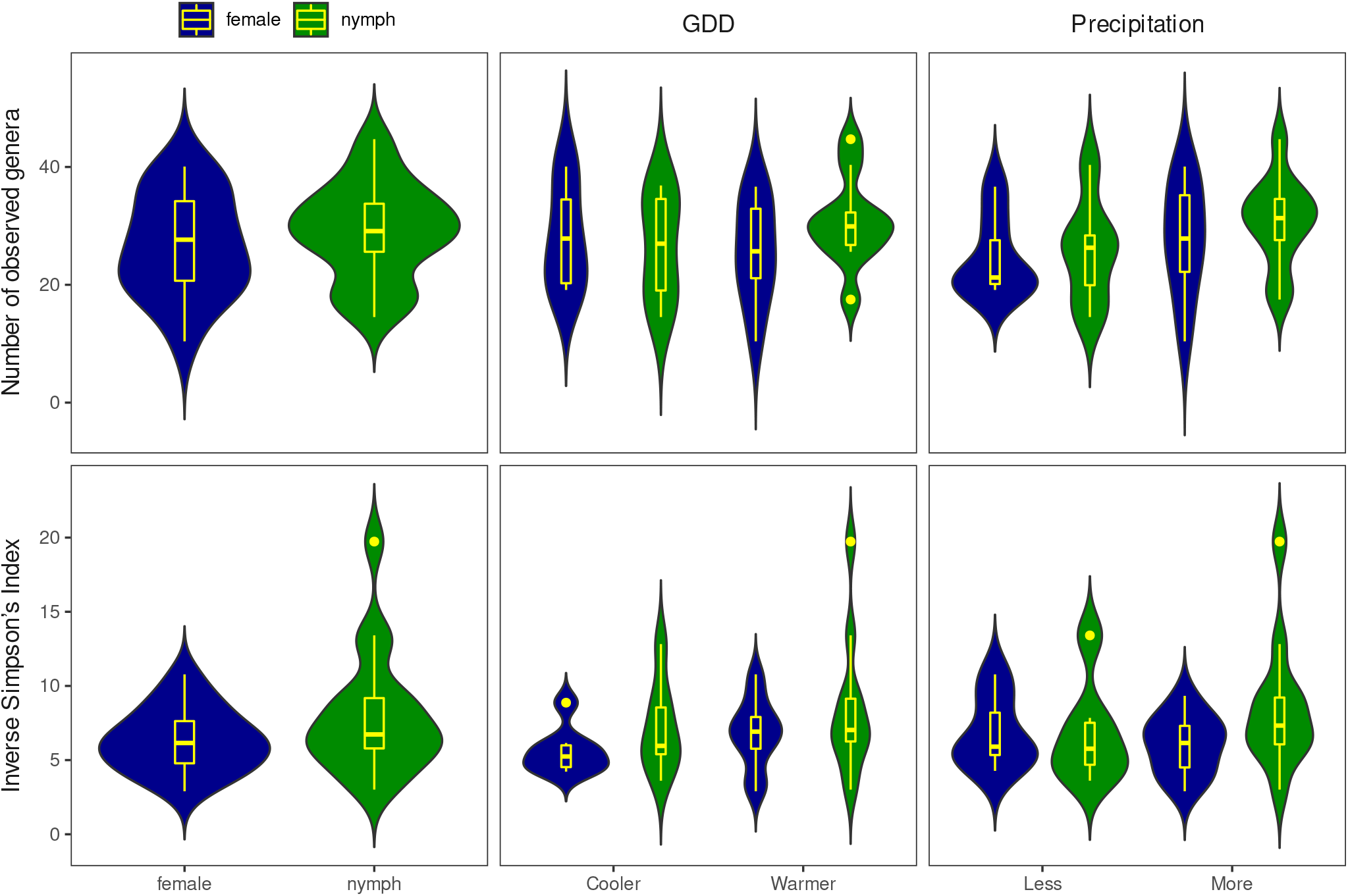
Richness and evenness of *Ixodes ricinus* bacteriome by sample groups. The numbers of observed species (richness) and the Inverse Simpson’s Index (evenness) as *α*-diversity metrics are presented in the form of a violin and box plot combination. These indices were calculated from 1000 iterations of rarefied OTU tables with a sequencing depth of 158. The average over the iterations was taken for each sampling site. The violin plot shows the probability density, while the box plot marks the outliers, median, and IQR.

### Core bacteriome

The core bacteriome is composed of the genera shown in Figure 3. For the list of species identified within genera see supplementary information.

**Figure 3.**
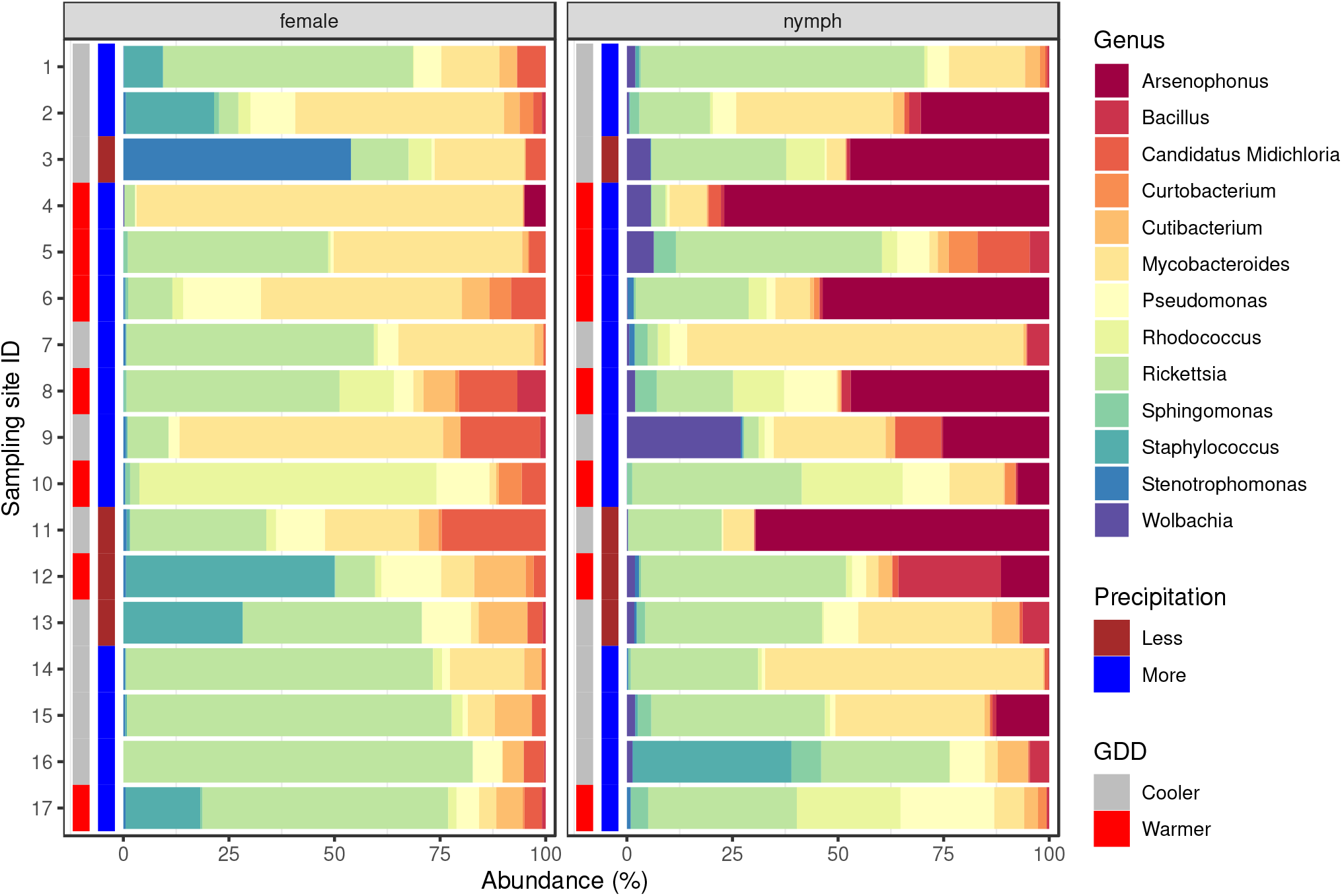
Core bacteriome composition of *Ixodes ricinus* samples. The relative abundance is plotted for the females and nymphs. Besides the bacterial genera of the core bacteriome, the environmental condition (growing degree-day (GDD) and precipitation) categories of sampling locations are also marked.

The variability of the samples’ core bacteriome genus profiles (*β* -diversity) is visualized by PCoA ordination (Fig 4) based on weighted UniFrac distance. The PERMANOVA analysis of the bacterial composition on the genus level revealed a significant (p=0.005) difference between the samples originating from females and nymphs. The core bacteriome of female samples showed no significant distance between either GDD (p=0.444) or precipitation (p=0.244) categories. Similarly, there was no significant difference between groups within nymphs (GDD p=0.108, precipitation p=0.722).

**Figure 4.**
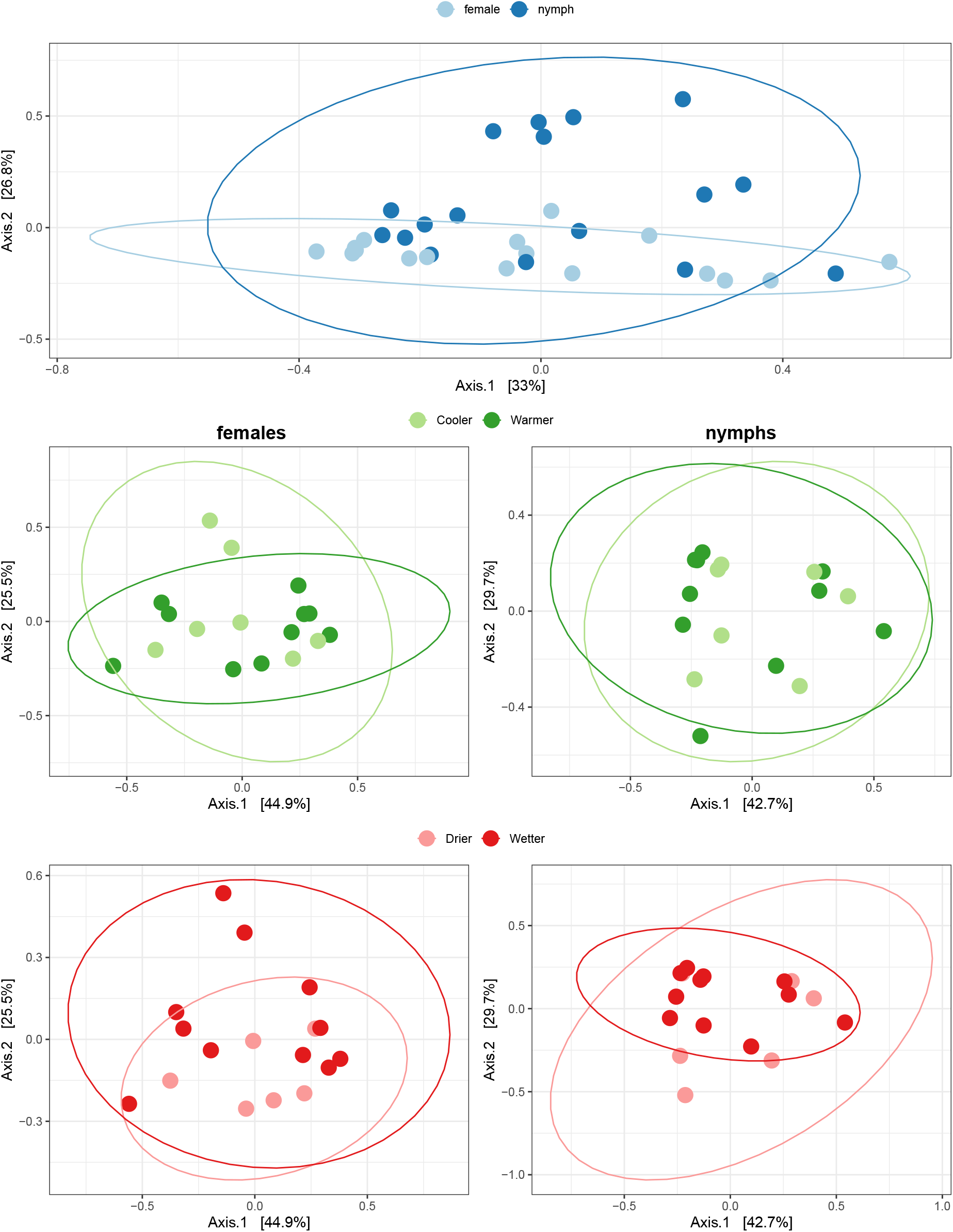
Principal coordinate analysis (PCoA) plots of *β* -diversity estimated based on the core bacteriome of *Ixodes ricinus* samples.

#### Abundance differences

The abundance differences (log2 median fold change, Log2FC) of groups per taxon comparison are summarized in Figure 5. Comparing females and nymphs, the following genera showed significant (adjusted p<0.05) differences in abundance: *Ar-senophonus, Bacillus, Candidatus Midichloria, Rhodococcus, Sphingomonas, Staphylococcus, Wolbachia*. In the female samples, according to GDD, the following were found differentiating: *Curtobacterium, Pseudomonas, Sphingomonas*. There was no genus with a significant difference in precipitation categories in females. In the nymphs, *Curtobacterium* showed a significant variation between GDD groups and *Bacillus* and *Curtobacterium* between precipitation levels.

**Figure 5.**
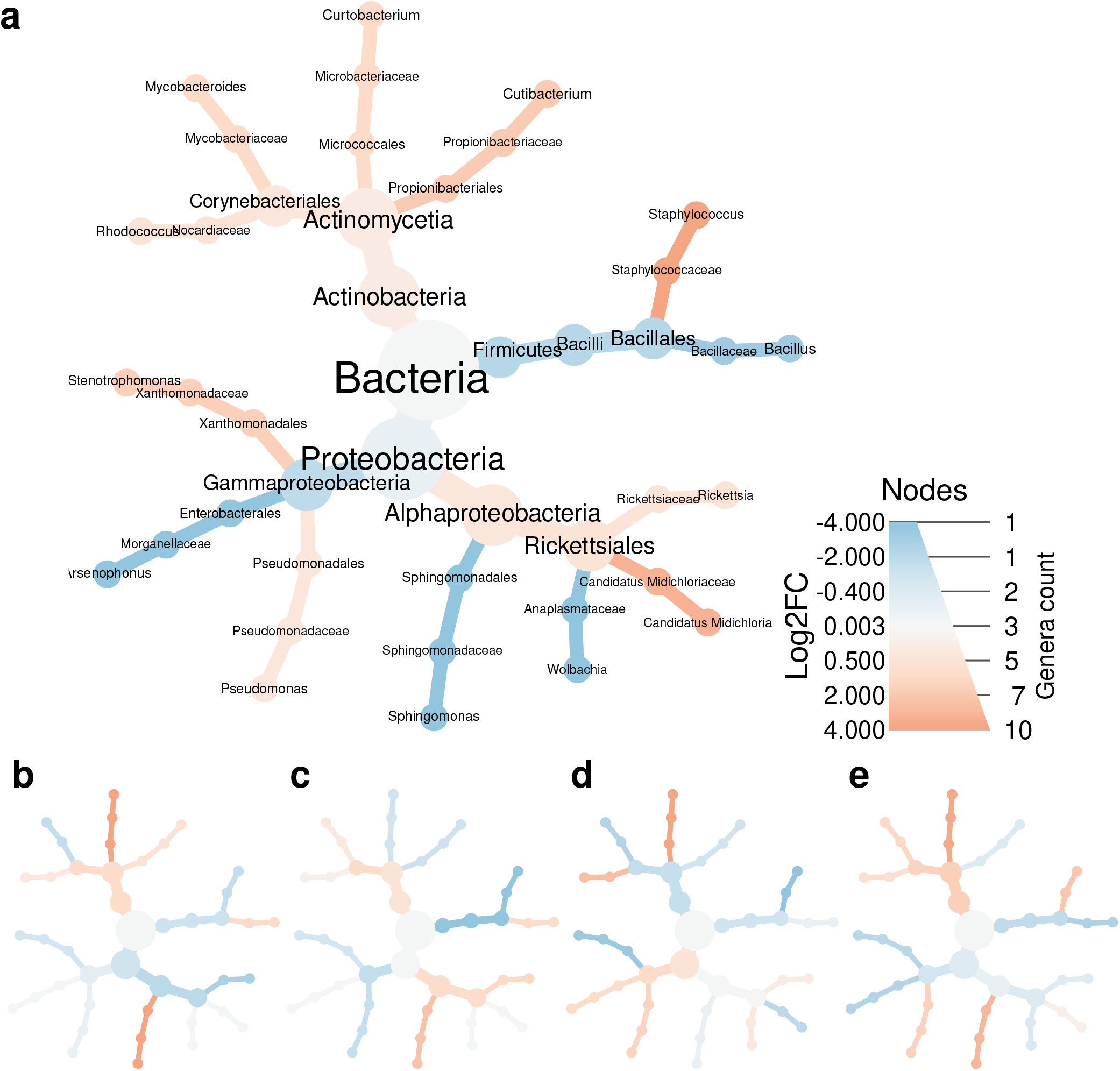
Core bacteriome abundance fold changes by taxonomic ranks. The colours represent the log2 fold change (Log2FC) of the median abundances of the compared groups. The subfigure **a** shows the ratio of the abundances in females comparing nymphs as a reference. Figure **b** compares female samples from warmer areas to cooler ones. Figure **c** compares samples from drier areas to those from wetter areas among females. Comparisons of GDD groups in nymphs are shown in figure **d**, while comparisons of precipitation groups are shown in figure **e**.

**Figure 6.**
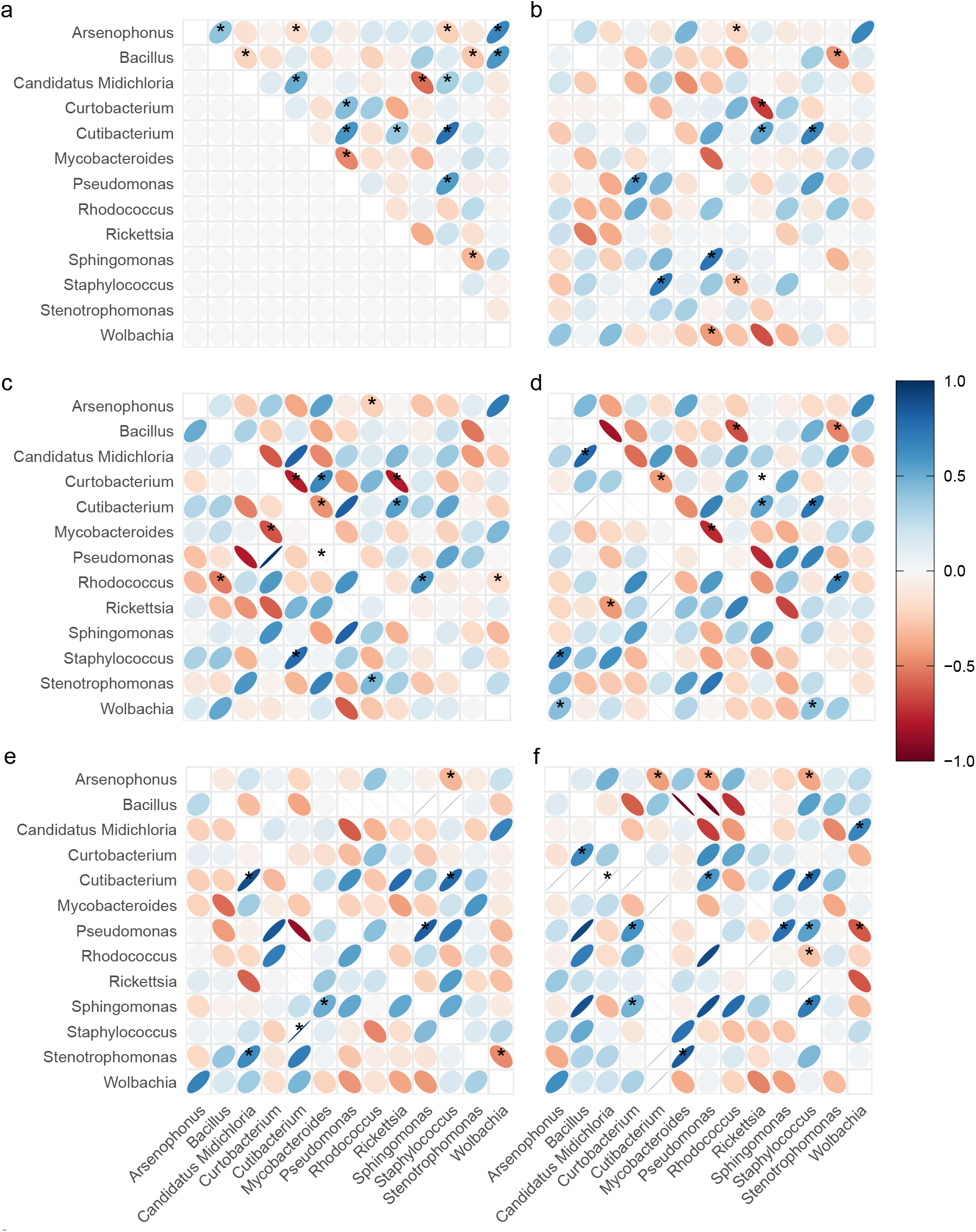
Bacteria genera abundance correlations. The correlation in all ticks is shown in Figure **a**. The lower half of figure **b** is obtained in nymphs and the upper half in females. In figure **c**, the correlations in females from cooler environments are in the lower, and those from warmer environments are in the upper triangle. In figure **d**, the correlations for females from drier environments are in the lower triangle and those from the wetter environment in the upper one. In figure **e**, the correlations of nymphs from cooler environments are in the lower triangle, and those from warmer environments are in the upper triangle. In figure **f**, the correlations of the nymphs from drier environments are in the lower triangle, and those from a wetter environment are in the upper triangle. Significant (p<0.05) relationships are marked by *.

## Discussion

Our shotgun sequencing-based microbiome analysis assesses an in-depth characterization of *I. ricinus* nymphs and adult females collected from a climatically representative set of sampling points in Hungary. Our study revealed a comprehensive picture of the bacterial diversity and associations in various host categories in *I. ricinus* ticks over the various climatic regions of Hungary.

While *I. ricinus* ticks had previously been linked with relatively higher alpha diversity scores than other common tick species,^34^ no significant difference was observed among the nymph and adult female stages or among warmer-colder and drier-wetter areas of origin. Carpi and colleagues found that the bacterial communities of geographically distant ticks of the same developmental stage differ more from those from the same regions.^35^ Moreover, Batool and colleagues found that in Ukraine, a neighbouring country to Hungary with an area around 6.5 times as big, the alpha diversity analyses demonstrated differences in tick microbiota patterns of various administrative regions.^36^ For the dissimilarity of the samples’ core bacteriome genus profiles (beta diversity), inter-regional comparison of developmental stages (females or nymphs) produced significant differences, while developmental stage-wise testing of the climatic condition-associated localization did not. Similarly, Batool and colleagues described that tick sex comparisons resulted in significant differences on various beta-diversity tests regardless of the area of origin.^36^ While these results are interesting to compare, the different study models and testing methods should be considered.

During the tick-collection phase of our study, three categories of ticks were evaluated for metagenome sequencing: nymphs, adult females, and adult males. Nevertheless, the sequencing of adult males was rejected as the number of males was lower than nymphs and adult females at all sampling points. Sex ratio shifts by ticks are not uncommon.^37,38^ The elucidation of the reason for the observed shift towards females in the adult life stage in our study would require further investigation. Nonetheless, the presence of certain maternally inherited genera, namely *Arsenophonus, Rickettsia, Spiroplasma* and *Wolbachia* in the metagenomes is noteworthy. These genera are described to induct parthenogenesis, feminize or kill males and, thus, manipulate the reproduction of their host species towards the production of daughters.^39,40^ Sex ratio skewness may be adaptive from the perspective of upper-generation ticks since it diminishes the competition of related males in tick-dense localities by reducing their numbers while increasing the numbers of related females that can be fertilized by the smaller male population.^41,42^

Besides the members of the core bacteriome of nymphs and adult females, reads deriving from further bacterial genera with relatively lower abundance rates but high clinical relevance (pathogens), such as *Anaplasma, Bartonella, Borrelia, Borreliella* and *Ehrlichia* have also been detected. While *Anaplasma phagocytophylum*, the cause of *Anaplasmosis* has previously been associated with the presence of a tick parasitoid, *Ixodiphagus hookeri*, based on its lifestyle and its mode of hunting,^43^ its positive correlation with the genera of *Arsenophonus* and *Wolbachia*, endosymbionts of *I. hookeri*, was not affirmed.^12^ Within our study, the geolocations where *Anaplasma* sp. occurred did not match the detection points *I. hookeri* in Hungary described in a recent study.^44^ Despite the high incidence rates of Lyme disease in Europe,^45^ *Borreliella burgdorferi* itself was not detected in our samples, while other species of *Borrelia* and *Borreliella* were identified. Since the causative agent of Lyme disease is normally mentioned as *B. burgdorferi* sensu lato,^46,47^ referring to this species broadly, involving other members of the genera as well, the presence of *B. garinii, B. valaisiana, B. afzelii, B. coriaceae* and *B. miyamotoi* can also be associated with this common disease. *Bartonella* spp. and *Ehrlichia* spp. are pathogens that are also often isolated in European settings.^48–50^ The presence of pathogens may influence the composition of tick microbiota.^51^

Some genera constituting the core bacteriome, namely *Arsenophonus, Candidatus Midichloria, Rickettsia* and *Wolbachia* are believed to be maternally inherited or strongly tick-associated due to direct or indirect reasons, while the members of *Bacillus, Curtobacterium, Cutibacterium, Mycobacteroides, Pseudomonas, Rhodococcus, Sphingomonas, Staphylococcus* and *Stenotrophomonas* are related to soil, water, plants, skin or mucosa of vertebrates, thus may rather be acquired from the environment of the ticks.^12,12,36,52,53^ The representatives of the environmental genera may either participate at ticks’ transient or long-term microbiota. Several environmental, tick gut or surface bacteria cause opportunistic infections in humans, especially in patients with an immunocompromised history.^54,55^ Even though certain bacteria, such as cutibacteria and staphylococci, may be considered as bacterial contaminants from the sample processing steps,^12^ according to a recent study, the effects of a possible, minor level contamination are not noticeable in the overall relative bacterial abundance.^36^ Nevertheless, it is possible that some environmental genera were only present at the cuticle of the collected ticks regardless of the repeated laboratory washing steps with 99.8% alcohol.

Except for *Cutibacterium, Mycobacteroides, Rickettsia* and *Stenotrophomonas*, genera constituting the core bacteriome showed statistically significant alterations in the examined tick groups differing in life stage and climatic condition-associated geographical origins. This finding is plausible considering the fact that both the life stage and the season-associated climatic conditions are very important in the composition of the tick bacteriome.^56,57^

The members of *Rickettsia* are maternally inherited or transstadially passed symbionts^58,59^ that may cause tick-borne infections^58^ and have been described to be dominant in tick microbiomes in several studies,^35,60–64^ However, high rickettsial abundance rates are not necessarily present in the ticks. Certain studies presented relatively low rickettsial genome fragment counts.^36^ The reason for these alterations is that the appearance rates of *Rickettsia* spp. vary between the geographical populations of ticks. According to a recent study, the members of *Rickettsia* are more abundant in ticks from forests,^53,65^ where ticks included in the present study also derived from. Considering that nymphal rickettsial counts were high in other studies as well,^60^ while male ticks are normally associated with relatively fewer members of the genus *Rickettsia*,^60,61^ the number of rickettsias seems to decrease throughout the life of male ticks gradually. This theory is in line with a study on microbiome changes during tick ontogeny.^66^ On the other hand, certain rickettsial species are also associated with male killing, which may also cause the decreasing numbers associated with male adult ticks.^67^ *R. helvetica* and *R. monacensis*, two species identified in our samples as well, have been connected with the presence of adult *I. hookeri* wasps that are the parasitoids of ticks. The reason for the association may either be related to the role of the parasitoid wasps in the circulation of rickettsias among ticks or with digested bacterial DNA in the wasp body lumen.^68^

Even though the members of Mycobacteroides are considered as either long-term or transient environmentally derived residents of the tick microbiota, recent studies revealed that certain species might be able to multiplicate inside the host and be transovarially inherited.^52,69^ Nevertheless, the relative abundance of the species seems dependent on the geographic region,^62^. In our study, a statistically non-significant shift could have been observed in the direction of cooler and dryer regions. Considering climatic tendencies, Thapa and colleagues reported higher mycobacterial genome fragment counts in ticks from Texas than from Massachusetts,^62^ which shows a contrary mycobacterial temperature preference. The observed differences may be caused by the different species-level *Mycobacterium* composition in the U.S. and in Europe. Since several *Mycobacterium* species may be pathogenic to humans or to animals, this finding may have a public health significance, especially in the light of cureent climate change trajectories.

*Cutibacterium* and *Stenotrophomonas* are common environmental bacteria that have been associated with ticks in several studies.^12,34,36^ *Cutibacterium* is strongly associated with ticks in forests,^65^ where our *I. ricinus* ticks are derived from. *Stenotrophomonas maltophilia*, a species identified in our samples, is an opportunistic pathogen often isolated from the infections of immunocompromised individuals.^55^

The majority of the genera constituting the core bacteriome were found to demonstrate significant differences among various life stage and climatic condition associated tick host groups. With the abundance rates of specifically tick-associated genera, *Arsenophonus, Candidatus Midichloria* and *Wolbachia*, the significant differences among adult female ticks and nymphs were clearly explainable.

*Arsenophonus*, which showed significantly lower abundance rates in females than in nymphs, is a widespread, mainly insect-associated bacterial genus with a wide spectrum of either parasitic or symbiotic host relations.^53,70^ While the high number of *Arsenophonus*, more precisely *A. nasonie* associated reads could suggest the dominance of this genus in the nymph microbiota; its presence is associated with *I. hookeri*, a parasitoid wasp of ticks that is supposed to have a wide geographical range worldwide. *I. hookeri* oviposits to larval and nymphal hard tick hosts. Its eggs can only develop in engorging or engorged nymphs.^71,72^ If the tick immune system-borne encapsulation of the *I. hookeri* eggs is not successful, the eggs hatch. Larvae start consuming the tick tissues and thus cause the death of the host.^71^ Furthermore, Bohacsova and colleagues found that *A. nasoniae*, an endosymbiont of encyrtid wasps identified in high numbers in our nymph samples, is only detectable in tick nymphs parasitized by the wasp.^68^ According to these findings, nymphs harboring *A. nasoniae* do not often reach the adult stage due to the parasitoid wasp, *I. hookeri*. Thus *A. nasoniae* deriving reads demolish from the adult tick population specifically due to the death of nymph hosts. While the developmental stage dependence of *I. hookeri* associated *A. nasoniae* is beyond question; the median abundance of this bacterial genus also showed differences in nymph groups of different climatical regions. Nymphs collected from warmer and rainier areas harbored more reads deriving from *Arsenophonus* spp. than ones collected from warmer areas. Attempts have been made to use *I. hookeri* as a means of biological control of ticks for approximately 100 years^72^ and the consideration of climatic conditions as underlying causes for unstable control technique success rates may improve current biological control methods. Moreover, *A. nasoniae* is described to be male-killing in several wasp species,^73^ but in *I. hookeri* adult wasp males were also infected by *A. nasoniae*, although the emergence ratio of males to females was 1:3.6 in the infected populations.^68^ Nevertheless, the presence of this bacterium may underlie decreased *I. hookeri* numbers at certain habitats or could even have contributed to insufficient abundance rates by biological tick control purposes, attempted mass releases of the parasitoid wasps in the past.^74,75^

The genus of *Wolbachia*, having a significant, strong abundance rate shift to nymphs is reported to share several characteristics with *Arsenophonus* that affect tick life and, potentially, tick population size. One of the identified species, *W. pipientis*, is strongly related to the presence of *I. hookeri*.^76,77^ Thus, the reason for the high number of wolbachial reads in nymphs may be the same as in the case of the *Arsenophonus* genus. *Wolbachia* spp. are also known to kill male hosts in some host insect species selectively,^39,78^ while in others it appears to be nonmale-killing.^79^ To the best of our knowledge, no studies exist on the effect of *Wolbachia* spp. on *I. hookeri*. Furthermore, similarly to the case of *Arsenophonus*, warmer and rainier areas had a slight positive effect on the number of wolbachial reads in nymphs although this effect was weaker than in the case of *Arsenophonus*. Due to the nymphal loads of reads from *Arsenophonus* and *Wolbachia*, the prevalence of *I. hookeri* in the samples and at the sampling points in Hungary can be strongly hypothesized. Evidence for the presence of these parasitoid wasps has recently been discovered in Hungary.^44^

By *Candidatus Midichloria*, the strong, statistically significant difference that was observed for females among adult female and nymph ticks is in agreement with the results of other research groups and has an explanation in the scientific literature. Studies exploring the inter-sex microbiome differences of adult *I. ricinus* ticks^34,36,80,81^ demonstrated much higher abundance rates of *Candidatus Midichloria* in females than in males independent to the regions of tick collection. The unique reason for this is that *Candidatus Midichloria mitochondrii* (CMM) invades the mitochondria of the cells within the ovaries of the female ticks. Despite the multiplication of the bacteria that consumes many ovarian mitochondria, the tick oocytes are expected to develop normally. CMM is described as being vertically transferred to all eggs.^80,82^ Even though the nymphal sex ratio of ticks may not be exactly 1:1,^83^ according to the available evidence, the presence of CMM does not result in sex ratio distortion in ticks.^80^ Simultaneously, it has been observed that CMM is transferred to both male and female larvae, but later, during the nymph stage, its specialization occurs toward females.^81^ Thus, a possible reason for the difference in CMM abundances among females and nymphs may be that many nymphs are males. Furthermore, the multiplication of CMM appears to increase after engorgement.^81^ Accordingly, the relatively low numbers of CMM might also be associated with the lack of engorgement among the nymphs collected for our study. Since tick collections occurred between the end of March and the middle of May, which represents the beginning of the activity period of nymphs,^84^ information on the overall engorgement status based on CMM counts appears to be realistic.

With the environmentally derived bacterial groups (*Bacillus, Curtobacterium, Pseudomonas, Rhodococcus, Sphingomonas, Staphylococcus*), the explanation of statistically significant life stage and climatic condition-wise differences in the bacterial composition appeared to be present.

*Bacilli* inhabit a broad range of environments, ranging from soils to insect guts and some species can be pathogens.^35^ The finding that the number of reads deriving from bacilli was significantly higher in nymphs than in adult females may originate from the choice to assess adult female ticks and exclude adult males from the study. While nymphs appeared in the study as a mixed population of males and females, the adult males collected at the sampling points were not represented due to their relatively low number compared to adult females. A study on the microbiome of *Rhipicephalus sanguineus* ticks describes a strong shift of *Bacillus* spp. towards the male tick population. Since our study only contained males in the nymph population, the observed shift may represent the male-relatedness to *Bacillus* spp. The nature of the relatedness of male ticks and *Bacillus* spp. has not yet been characterized.^60^ Nonetheless, *Bacillus* spp. are described not to be present in every tick microbiota^62,85^ and the detection rates of the members of this genus appear to show significant regional differences.^36^ Within our study, reads deriving from *Bacillus* spp. showed significantly higher abundances in nymphs from areas with more precipitation. Furthermore, Fernández-Ruvalcaba and colleagues reported that *B. thuringiensis* strains significantly elevated mortality and demolished oviposition and egg hatch among adult, pesticide-resistant *Rhipicephalus microplus* ticks.^86^ Moreover, certain strains of *B. wiedmannii* are known, while *B. thuringiensis* strains are even commercialized as insecticides or nematicides for biological pest control,^87,88^ although the entomopathogenic effect of environmental strains proved to be more present.^89^ Both above-mentioned *Bacillus* species were present in our samples, which can decrease *I. ricinus* numbers, although potentially in both age groups. The key to the biopesticide characteristics of these bacilli is the formation of functional crystalline *Cry* proteins that are specific toxins. Other entomopathogenic bacteria such as *B. cereus*, also present within our samples, are opportunistic as they act in secondary, non-specific pathways facilitated by certain weakening motives, such as the presence of *Cry* proteins.^88^ Thus, the shift in *Bacillus* abundance towards nymphs rather appears to occur due to the sex determination of the adult ticks included in the study.

Although within our samples, significantly more *Rhodococcus* reads are derived from adults than from nymphs, this result may be based on the following factor. The dominating species, *R. fascians*, is a common bacterial phytopathogen that interacts with a broad array of plants, causing their malformation.^90^ Older ticks may have had more opportunities for encounters with these environmental bacteria than younger ones. Higher *Rhodococcus* read counts in adults align with René-Martellet and colleagues’ findings.^60^

Association with *Staphylococcus* spp. that were significantly more abundant in adult females than in nymphs may be explained with similar reasoning. *Staphylococci* are common findings related to ticks^52,61^ that often appear on the skin and mucous membranes of the hosts of the ticks as well.^52^. Ticks carrying staphylococci may have already engorged and thus encountered these bacteria. Considering that the number of engorged nymphs appeared to be relatively low within our samples according to the reasons explained by the genus *Candidatus Midichloria*, fewer opportunities to encounter the members of this genus by nymphs are possilbe. Moreover, the relative abundance of *Staphylococcus* spp. also appears to be dependent on the region of tick collection^62^ and, as described earlier, staphylococcal hits may also derive from contamination.^36^

The finding that the members of another environmental bacterial genus, *Sphingomonas*^12^ were identified with significantly higher abundance rates in nymphs than in adults is controversial to the hypotheses of *Rhodoccus* spp., *Staphylococcus* spp. and to the related findings of other authors.^66^ Another finding was that among adult females, the abundance rate of *Sphingomonas* spp. was significantly higher in warmer sampling areas than in cooler localities, while by nymphs, the temperature-wise difference was not relevant. Interestingly, another research group found that *Sphingomonas* spp. were much more abundant in adult ticks kept at 4 °C, than by those at 20 °C, 30 °C or at 37 °C. Moreover, according to their study, *Sphingomonas* was among the most abundant bacterial genera at 4 °C by males.^61^ The reason for this discrepancy with our results on temperature-wise abundance may rest on the species-level *Sphingomonas* composition. Additionally, interactions with other bacterial groups may also underlie our findings. In any case, further studies would be required to elucidate the findings of *Sphingomonas* populations and confirm or invalidate our hypotheses.

Statistically significant climatic condition-wise alterations in adult female and nymphal stage ticks only appeared in environmental bacterial genera, namely *Curtobacterium* and *Pseudomonas*. While the former appeared significantly more abundantly in warmer environments both for adult females and nymphs, a significant preference for little precipitation was observable by nymphs. This finding may be relevant in light of climate change, mostly because of a dominant species, *C. flaccumfaciens* which is a phytopathogenic bacteria with economic significance^91^ and has also been isolated from a child with fatal septicemia.^92^

Reads from the genus of *Pseudomonas* were detected with significantly greater abundances at areas with higher GDD in adult females, while temperatures appeared not to influence the relative abundance of *Pseudomonas* spp. in nymphs. The formation of biofilms might explain this finding as some *Pseudomonas* strains have better biofilm formation capacities at warmer temperatures.^93,94^ Thus, adult ticks from warmer environments that have already survived at least one summer, according to our knowledge of tick life cycle, may harbour more bacteria from the *Pseudomonas* genus that managed to form biofilms and so became more steadily present. According to a study, the egg wax composition of the cattle tick, *Rhipicephalus microplus* is able to inhibit the biofilm formation of *P. aeruginosa*.^95^ Thus, certain mechanisms may influence the abundance of *Pseudomonas* species at earlier tick life stages. The exact composition of *Pseudomonas* strains should be examined to evaluate this hypothesis as the genus is very versatile. At the same time, *Pseudomonas* spp. have been related to both nymphs and adults in other publications,^36^ while certain studies report more stable *Pseudomonas* spp. appearance rates in males than in females.^34,62^

All the above-mentioned bacterial groups share another factor, namely the interaction among the bacterial community members, this may be responsible for the microbial pattern assessed. In order to elucidate the possible influence of certain bacterial co-occurrences, the correlation analysis of the bacterial genera was performed in each development stage and climatic category. Positive correlations among the taxa of microbial communities may be interpreted as the reflection of shared habitat or environmental condition preferences, cooperative activities, such as cross-feeding^96^, or the representation of functional guilds performing complementary or similar functions.^97^ In contrast, negative correlations may indicate competition for limiting resources, niche partitioning, inequivalent resistance to losses or active negative interactions.^96,97^ In contrast to other studies,^12^, the number of positive and negative correlations was balanced within our samples.

All samples considered, *Arsenophonus* and *Wolbachia* correlated positively, which is assumed to have occurred due to the presence of *I. hookeri*. Furthermore, *Rickettsia* was observed to be negatively correlated with *Curtobacterium* spp. in several development stages and climatic groups, which may indicate positive public health consequences and a possible step towards a future tick control tool. In contrast, *Candidatus Midichloria* spp., that were previously detected in positive correlation with *Rickettsia* spp.^12^ showed no correlation in our samples. Juxtaposing another publication in the field,^12^ neither the *Pseudomonas* and *Rickettsia*, nor the *Bacillus* and *Rickettsia* pair showed any correlation. Correlations among environmental bacteria of various climatic categories are likely based on their similar environmental preferences and the previously mentioned interaction types. Due to the seasonal variability of environmental bacteria, the number of tick-borne, potentially pathogenic bacteria may increase or decrease correspondingly with the taxa of environmental origins.^12^ Accordingly, the presence or absence of certain environmental taxa may reflect the temporal dynamics of certain pathogens. The public health significance of this finding is particularly significant in light of climate change and potentially varying climatic conditions around the globe. Nevertheless, the interpretation of co-occurrence patterns and the nature of these correlations require further studies as abundance shifts depend on multiple factors. Thus, bacterial and parasitological interconnections are not exclusively responsible for the variations. Moreover, our results were obtained from the sequencing of entire tick individuals and thus lacked finer, e.g., organ scale considerations of the correlating taxa.

## Conclusion

Here we reported the identification of the *I. ricinus* bacteriome-associated findings in adult females and nymphs collected from a climatically representative set of sampling points in Hungary. These results allowed us to show that (1) the *I. ricinus* bacteriome is dependent on the temperature and precipitation history of the geolocation of sampling; (2) the *I. ricinus* bacteriome is not stable in the developmental stages of the ticks; (3) based on the bacteriome patterns identified, the identified developmental stage-wise alterations may be associated with the presence of certain tick parasitoids that exclude the option of reaching the adult age; (4) tick-borne disease pathogens are widely distributed at the climatically representative sampling points.

In the future, developmental stage and climate-associated microbial differences and correlations identified in this ecosystem study could be confirmed with experimental approaches and complemented with further metagenome studies to achieve sufficient data volumes of tick microbial inter-relatedness and exploit them as promising resources for novel tick control strategies.

## Declarations

### Ethics approval and consent to participate

Not applicable.

### Consent for publication

Not applicable.

### Availability of data and material

The short-read data of samples are publicly available and accessible through the PRJNA828115 from the NCBI Sequence Read Archive (SRA).

### Author contributions statement

NS takes responsibility for the integrity of the data and the accuracy of the data analysis. NS and RF conceived the concept of the study. EK, GM, and MG performed sample collection and procedures. AGT, MP, and NS participated in the bioinformatic and statistical analysis. AGT, HY, LM, and NS participated in the drafting of the manuscript. AGT, GM, NS and OK carried out the manuscript’s critical revision for important intellectual content. All authors read and approved the final manuscript.

## Acknowledgements

Not applicable.

## Authors’ information

Not provided.

## Supplementary information

### Species in core bacteriome

*Arsenophonus endosymbiont of Aphis craccivora, A. endosymbiont of Apis mellifera, A. nasoniae*.

*Bacillus frigoritolerans, B. cereus, B. mycoides, Bacillus* sp. 7D3, *B. thuringiensis, B. wiedmannii*.

*Candidatus Midichloria mitochondrii*.

*Curtobacterium flaccumfaciens, Curtobacterium* sp. 24E2, *Curtobacterium* sp. BH-2-1-1, *Curtobacterium* sp. C1, *Curtobacterium* sp. TC1.

*Cutibacterium acnes, C. avidum, C. granulosum, C. modestum*.

*Mycobacteroides [Mycobacterium] stephanolepidis, M. abscessus, M. chelonae, M. immunogenum, M. salmoniphilum, M. saopaulense*.

*Pseudomonas aeruginosa, P. alcaligenes, P. antarctica, P. azotoformans, P. brenneri, P. cichorii, P. congelans, P. eucalypticola*,

*P. extremorientalis, P. fluorescens, P. frederiksbergensis, P. graminis, P. lurida, P. moraviensis, P. orientalis, P. poae, P. prosekii, P. putida, P. qingdaonensis, P. rhizosphaerae, P. soli, Pseudomonas* sp. 15A4, *Pseudomonas* sp. CIP-10, *Pseudomonas* sp. DG56-2, *Pseudomonas* sp. HN2, *Pseudomonas* sp. HN8-3, *Pseudomonas* sp. LBUM920, *Pseudomonas* sp. NS1(2017), *Pseudomonas* sp. OE 28.3, *Pseudomonas* sp. S49, *P. stutzeri, P. synxantha, P. syringae, P. tensinigenes, P. trivialis, P. umsongensis*.

*Rhodococcus erythropolis, R. fascians, R. globerulus, R. koreensis, R. opacus, R. qingshengii, Rhodococcus* sp. AQ5-07, *Rhodococcus* sp. B7740, *Rhodococcus* sp. H-CA8f, *Rhodococcus* sp. MTM3W5.2, *Rhodococcus* sp. P1Y, *Rhodococcus* sp. PBTS 1, *Rhodococcus* sp. YL-1. *R. amblyommatis, R. asiatica, R. bellii, R. conorii, R. endosymbiont of Ixodes scapularis, R. helvetica, R. monacensis, R. parkeri, R. rhipicephali*.

*Sphingomonas aliaeris, S. alpina, S. insulae, S. koreensis, S. melonis, S. paucimobilis, S. sanguinis, S. sanxanigenens, Sphingomonas* sp. AAP5, *Sphingomonas* sp. FARSPH, *Sphingomonas* sp. HMP9, *Sphingomonas* sp. LK11, *Sphingomonas* sp. LM7, *Sphingomonas* sp. NIC1, *Sphingomonas* sp. PAMC26645, *S. taxi*.

*Staphylococcus arlettae, S. aureus, S. auricularis, S. capitis, S. epidermidis, S. haemolyticus, S. hominis, S. pasteuri, S. saccharolyticus, S. warneri*.

*Stenotrophomonas acidaminiphila, S*. indicatrix, *S*. maltophilia, *S*. rhizophila, *Stenotrophomonas* sp. 169, *Stenotrophomonas* sp. DR822, *Stenotrophomonas* sp. LM091, *Stenotrophomonas* sp. NA06056, *Stenotrophomonas* sp. SI-NJAU-1, *Stenotrophomonas* sp. SXG-1.

*Wolbachia endosymbiont of Anopheles demeilloni, W. endosymbiont of Ceratosolen solmsi, W. endosymbiont of Chrysomya megacephala, W. endosymbiont of Corcyra cephalonica, W. endosymbiont of Delia radicum, W. endosymbiont of Drosophila simulans, W. endosymbiont of Wiebesia pumilae, W. pipientis*.

### Pathogens

Taxon classification of short reads with Kraken2 resulted in hits for the following pathogenic bacteria. *Anaplasma phagocytophilum* was found in nymphs from sampling sites 3 and 7. Reads from *Borrelia coriaceae* were identified in females collected from sampling site 1. *B. miyamotoi* was found in females from sampling sites 2 and 16 and in nymphs collected from sampling sites 1 and 10. *Borreliella garinii* was at sampling site 11 in the nymphs. *B. valaisiana* was found in nymphs collected at sampling site 8 and 11. *B. afzelii* was found at sampling site 2 in nymphs. *Ehrlichia muris* related reads were found in female 12 and nymph 14 samples. Reads originating from the genus *Rickettsia* were found in all samples: *R. amblyommatis* in females (sample site: 5) and nymphs (sample site: 6); *R. asiatica* in females (1, 7, 12) and nymphs (1, 2, 10, 15); *R. bellii* in females (5) and nymphs (6, 10); *R. conorii* in females (8); *R. helvetica* in females (1, 3-5, 7-10, 13-17) and, nymphs (1-6, 8, 10, 11, 13-17); *R. monacensis* in females (3, 5, 8) and nymphs (1, 6, 10, 12); *R. parkeri* in nymphs (10); *R. rhipicephali* in females (5, 8) and nymphs (1, 6, 10). No species from the genera *Bartonella, Coxiella* and *Francisella* was found.

The result of the BLAST^26^ based taxon classification of the assembly-generated contigs is as follows: *Candidatus Odyssella thessalonicensis* L13 was found in females (sample site: 5); *Candidatus Rickettsia colombianensi* in females (5, 8, 15) and nymphs (4, 6, 10, 13); *Orientia tsutsugamushi* in females (5) and nymphs (10); *Rickettsia akari* in nymphs (6, 10); *R. asembonensis* in females (5, 8, 17) and nymphs (6, 8, 10, 13); *R. asiatica* in females (3, 4, 5, 7, 8, 14, 15, 16, 17) and nymphs (1, 2, 3, 4, 6, 8, 10, 11, 13, 14, 15, 16, 17); *R. australis* in females (5, 8, 15) and nymphs (3, 4, 6, 10, 11, 14); *R. bellii* in females (8) and nymphs (1, 6, 10); *R. canadensis* in females (3, 5, 7, 8, 14, 17) and nymphs (1, 3, 6, 10, 11, 15); *R. conorii* in females (5) and nymphs (6, 10, 15); *R. felis* in females (1, 5, 7, 8, 13, 15, 17) and nymphs (1, 3, 4, 6, 10, 11, 14, 15); *R. fournieri* in females (4, 5, 8, 15) and nymphs (1, 4, 5, 6, 10, 11, 15); *R. gravesii* in females (5, 8) and nymphs (3, 6, 10); *R. helvetica* in females (1, 3, 4, 5, 7, 8, 13, 14, 15, 17) and nymphs (1, 2, 3, 6, 8, 10, 11, 13, 14, 15, 16); *R. honei* in females (5, 7, 8, 14, 17) and nymphs (3, 6, 10); *R. hoogstraalii* in females (4, 5, 8, 15, 17) and nymphs (1, 3, 6, 10, 15); *R. japonica* in females (5) and nymphs (6, 10); *R. monacensis* in females (5, 8, 15) and nymphs (1, 4, 6, 10); *R. prowazekii* in females (5, 8, 14, 15) and nymphs (1, 6, 10, 11); *R. rhipicephali* in females (1, 3, 5, 8) and nymphs (1, 3, 6, 10, 11); *R. rickettsii* in females (5) and nymphs (6, 10); *R. sibirica* in females (5) and nymphs (6); *R. slovaca* in females (1) and nymphs (6, 10, 15); *Rickettsia* sp. MEAM1 in females (5, 7, 8, 15) and nymphs (6, 10, 11); *R. tamurae* in females (4, 5, 7, 8, 15, 17) and nymphs (1, 4, 6, 10, 13, 14, 15); *R. tillamookensis* in females (4, 5, 17) and nymphs (1, 2, 3, 6, 10, 11, 15); *R. typhi* in females (8) and nymphs (4, 6, 10); *Spiroplasma endosymbiont of Danaus chrysippus* in females (13).

